# Quantitative high-resolution imaging of mouse nephron formation to study Wnt signaling dynamics

**DOI:** 10.1101/2025.06.30.662340

**Authors:** Nobuko Tsuchida-Straeten, Simon Hammer, Aliaksandr Halavatyi, Christian Tischer, Gislene Pereira, Matias Simons

## Abstract

**Introduction:** Nephron formation is initiated when Wnt9b from the ureteric bud acts on nephron progenitor cells in the metanephric mesenchyme. For mouse embryonic kidneys, this process can be studied in real time using *ex vivo* organ cultures. Previous imaging methods relied on Transwell filters with a long distance between objective and sample as well as low signal-to-noise ratio due to the filter membrane. Moreover, Wnt signaling was previously visualized with the expression reporters *TCF/Lef:H2B-GFP*.

**Methods:** We developed an *ex vivo* culture system based on low-volume media and fibronectin coating for real time imaging. We further made use of endogenously tagged *Cherry-Lef1* and *Ctnnb1-Venus* reporter mouse lines as reporters for Wnt signaling during kidney development. Furthermore, we established an adaptive feedback microscopy pipeline to track the signal with high resolution.

**Results:** Using these approaches, we found that Cherry-Lef1 proteins are only expressed in the nascent nephron and derivatives. Expression is graded along distal-proximal nephron axis suggesting that the ureteric bud-derived Wnt9b is controlling its expression. Interestingly, we observed that Cherry-Lef1 gradient shifts from distal to proximal axis during nephron patterning. β-catenin*-*Venus, on the other hand, first became visible in the ureteric bud, then in the nascent nephron after Lef1.

**Conclusions:** Using a novel imaging approach we demonstrate a dynamic regulation of Wnt signaling that correlates with nephrogenesis in mouse kidney development. Combined with our two new Wnt signaling reporters, nephron formation can be studied in real-time with high resolution.

## Introduction

The Wnt/β-catenin pathway is known to play an important role to trigger the earliest stage of nephron formation. While nephron progenitor cells maintain a low level of Wnt signaling for self-renewal, high levels of Wnt signaling, induced by ureteric bud-derived Wnt9b at the branch corners, elicits a nephrogenic program, starting with conversion of the mesenchymal cells into the epithelial renal vesicle ^1-4^. For this mesenchymal-to-epithelial transition (MET), another Wnt, Wnt4, is needed which is expressed by the cells undergoing MET under the control of Wnt9b ^5,6^. Subsequently, the renal vesicles (RV) grow in the distal-proximal direction while differentiating into different nephron segments. The spatial and temporal regulation of Wnt activity is finely tuned by a network of antagonists, co-receptors, and intracellular modulators, ensuring proper kidney patterning and size ^7^. Aberrations in Wnt signaling components can result in a range of developmental disorders, including renal hypoplasia, dysplasia, cystic kidney disease ^8^ and consequences of low nephron number including chronic kidney disease in adults ^9,10^.

Early nephrogenesis can be studied in *ex vivo* kidney culture systems, for which several protocols exist ^11,12^. The most commonly used method is Trowell-type air-liquid interface culture ^13^ based on the use of commercial transwell membrane insert ^14^. However, this method requires long focal length between the specimen and objective, making it difficult to quantify images over time. Also, the signal-to-noise ratio is typically low as the laser has to pass through the membrane. Hence, this method is not ideal for live imaging. To monitor Wnt signaling during live imaging, so far only the *TCF/Lef::H2B-GFP* reporter has been used in embryonic kidney development ^15,16^.

In this study, we analyzed Wnt signaling dynamics during kidney development using two newly established endogenously tagged knock-in mouse lines, *Cherry-Lef1* and *Ctnnb1-Venus*, established in the Aulehla lab (EMBL, detailed report published elsewhere). Live imaging is performed on E12.5/13.5 murine embryonic kidneys with an *ex vivo* culture system that is based on a low-volume culture using fibronectin coating (**Fig. 1a, b**). While allowing normal growth of embryonic kidneys, this method is easy-to-handle and applicable for quantitative high-resolution microscopy. Visualization of Wnt signaling dynamics with these reporters allows the spatiotemporal analysis of nephron formation with unprecedented resolution.

**Figure 1.**
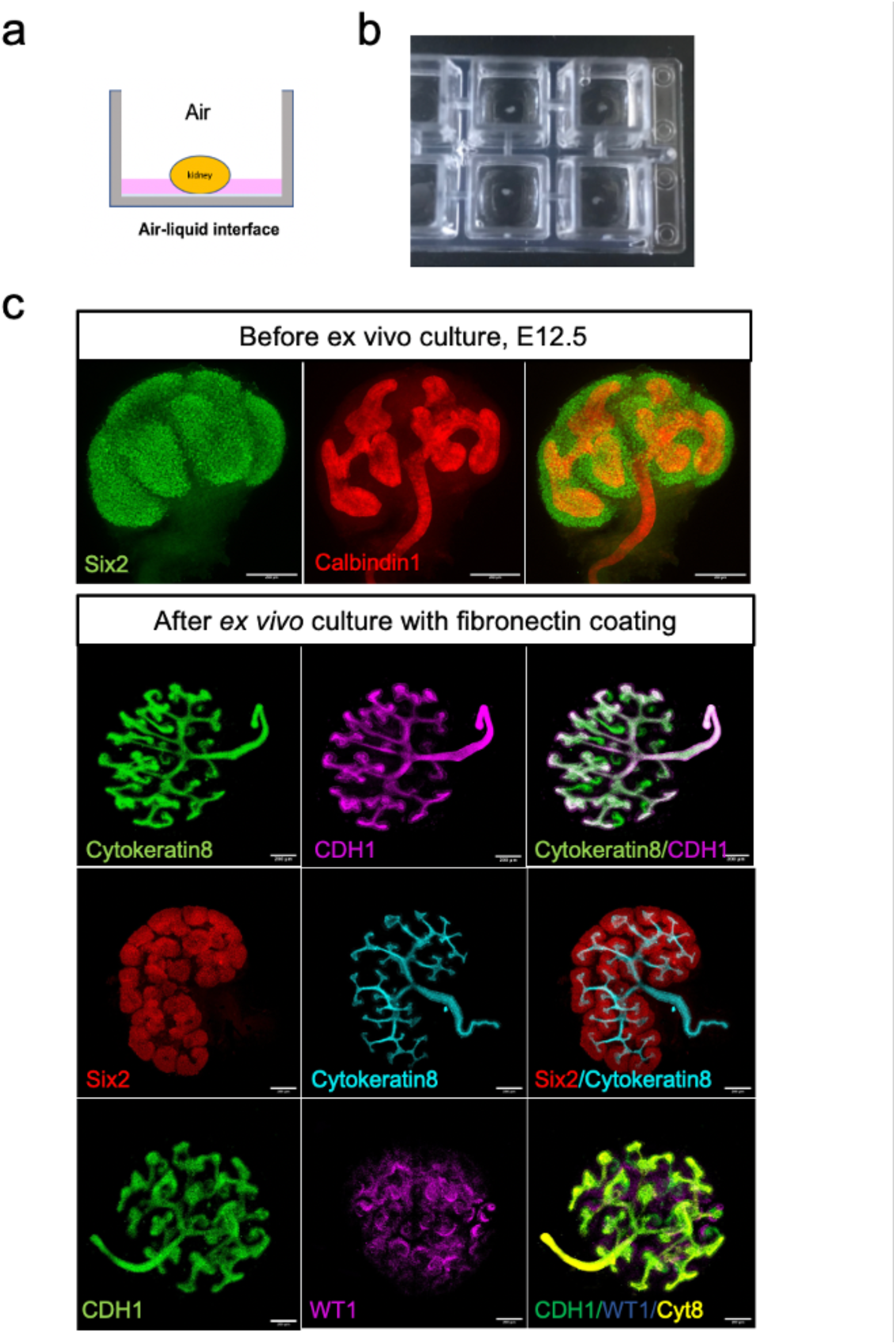
Development of mouse embryonic kidneys in fibronectin-coated glass slide chamber. **(a)** Schematic of *ex vivo* culture based on low-volume media in the chamber. **(b)** Picture of *ex vivo* system from above. **(c)** Whole mount staining of embryonic kidneys before and after *ex vivo* culture for 18 hrs using Six2, Calbindin1, Cytokeratin8, CDH1 and WT1 antibodies. Scale bar, 200µm,

## Results

### An *ex vivo* culture system with fibronectin coating for live imaging

Low-volume cultures utilizing silicone rings and glass cover slip have successfully been used for embryonic kidney organ culture ^11^. We adapted this method by cultivating E12.5 or E13.5 embryonic kidneys on a fibronectin-coated 8-well chamber slide in low-medium volume creating an air-liquid interface (**Fig. 1a, b**). By avoiding the membrane filter, signal-to-noise ratio is improved. Moreover, this method allows easy handling and can be used for imaging at multiple positions because the kidney samples are positionally fixed by the fibronectin coating and the microscope stage only has to cover short distances.

To evaluate kidney development with this method, we performed whole mount staining after 18 hours of *ex vivo* culture (**Fig. 1c**). While cytokeratin-8-positive ureteric buds showed adequate branching, Six2-positive nephron progenitor cell populations were localized around the ureteric bud tips. Furthermore, we observed a proper proximal-to-distal nephron patterning when staining for the podocyte marker WT1 and the distal tubule marker CDH1. When comparing developmental progression to the conventional Trowell method using commercial transwell membrane inserts, we could not find any differences with regard to the morphology and marker expression (**Fig. 1 - Supplement1**). This suggests that our fibronectin coating method allows for *ex vivo* kidney organ culture with normal development but improved handling and optical characteristics compared to the conventional method.

### Endogenous Cherry-Lef1 is expressed in the nascent nephron

Nephron induction and MET occurs at the branch corners of the ureteric tree where the concentration of Wnt9b secreted from the ureteric tree is highest ^17^. This ensures that a single nephron can connect with each branchpoint. In the nascent nephron, high β-catenin activity allows for TCF/LEF-dependent gene expression ^18^. To study spatial dynamics of Lef1, we generated a knock-in mouse line in which *mCherry* is inserted after the start codon in the endogenous *Lef1* locus (**Fig. 2a**). We observed that homozygous knock-in mice were viable and fertile, suggesting that the Cherry-Lef1 fusion protein is functional. We performed live imaging with this mouse line and detected the highest expression at the distal end of the nascent nephron (or RV; **Fig. 2b, c**) where the Wnt9b concentration is presumably the highest. Accordingly, the signal gradually decreased towards the proximal end (**Fig. 2d; Movie1; Movie4; Fig. 3c, d**). Whole mount staining with an anti-Cherry antibody shows that Cherry-Lef1 expression is observed in the nucleus and shows a specific spatial pattern in the nephron (**Fig. 2 - Supplement 1a, b, c**). Interestingly, we found that the Lef1 expression pattern has a conical shape with the narrow end being distal (**Fig. 2 - Supplement 1a, b, c**). This conical shape pattern was particularly obvious at E13.5 (**Fig. 2 - Supplement 1b, c**). In the ureteric bud, Cherry-Lef1 expression was absent unlike the previously used *TCF/Lef::H2B-GFP* reporter ^16^. These data suggest that endogenous Cherry-Lef1 is specifically expressed in the developing nephron forming a gradient along the distal to proximal axis.

**Figure 2.**
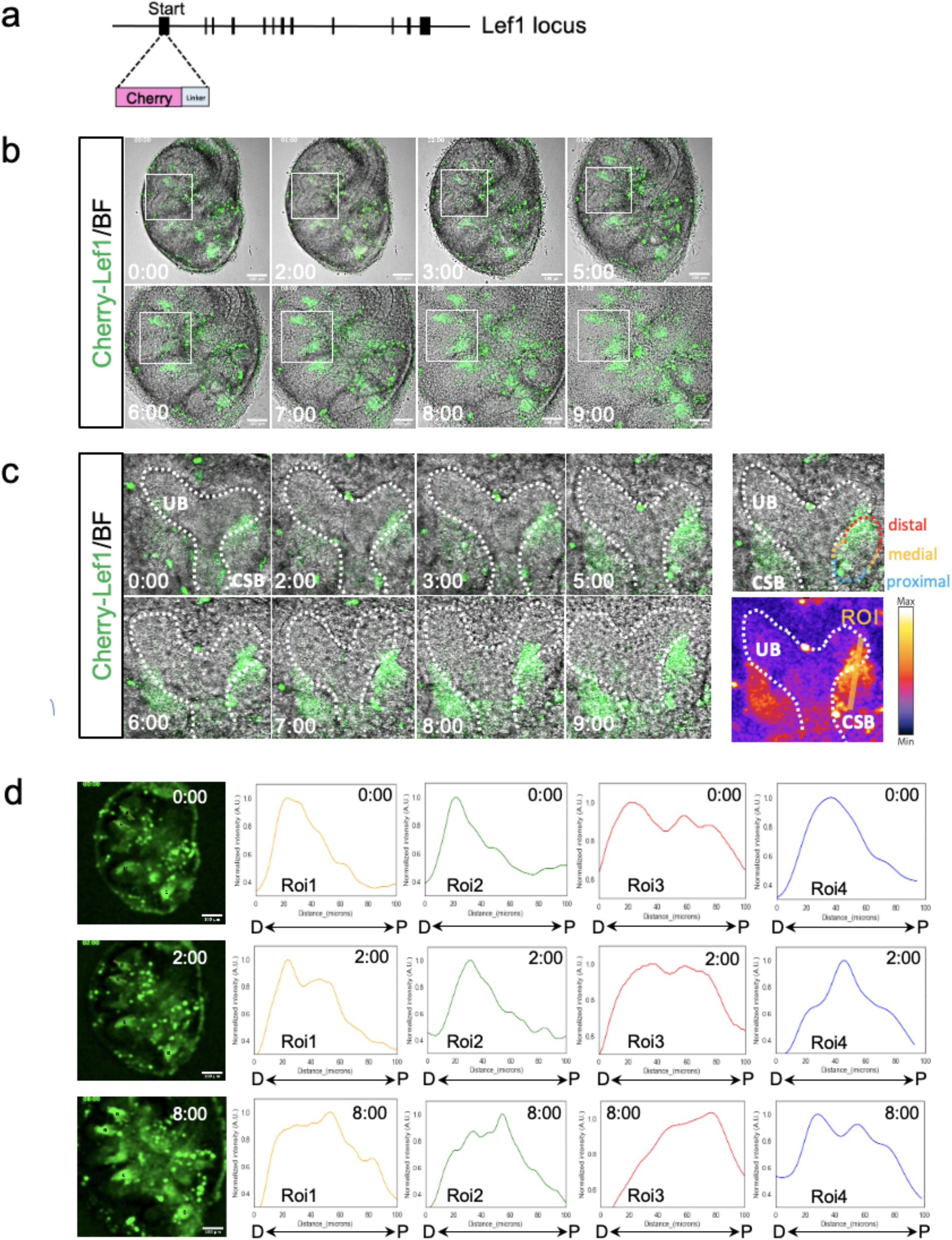
Generation of Cherry-Lef1 mouse lines to study Lef1 dynamics in the embryonic kidney. **(a)** Schematic representation of the tagging strategy for the Cherry-Lef1 mouse line. Lef1 was endogenously tagged with Cherry-linker. **(b)** Snapshot images of real time imaging at different time points(0, 2, 3, 5, 6, 7, 8 and 9hrs) in embryonic kidneys(E12.5) using Cherry-Lef1 mouse line. Scale bars,100µm. **(c)** Enlarged images of nephrons of (b). Roi was indicated as the orange line for quantification. **(d)** Quantification of Cherry-Lef1 signal intensity in nephrons. Left panel: Snapshot images at 0, 2 and 8hrs and roi(1-4) in 4 nephrons. Scale bars,100µm. The plots show normalized intensity with different roi(1-4) and time points(0, 2 and 8hrs). D: distal, P: proximal.

### Cherry-Lef1 gradient shifts along the nephron axis

β-catenin is the core effector of Wnt signaling that exists in two cellular pools. The cytoplasmic/nuclear pool acts as a transcriptional co-activator of TCF/LEF target genes, while the membrane-bound pool is a critical component of cadherin-based adherens junctions ^19,20^. To study this Wnt signaling marker, we generated a *Ctnnb1-Venus* knock-in mouse line where m*Venus* is endogenously tagged in the C terminal end of *Ctnnb1* locus. Homozygous mice were lethal but heterozygous mice were viable and fertile. We conducted real time imaging with this mouse line and found that the expression of β-catenin-Venus was highly dynamic during kidney development (**Movie2**). It first appeared in the ureteric tree and then after 4 hours at the distal end of the RV before spreading to the proximal end. In the ureteric bud and nephron epithelial cells, the signal mainly came from the apical junctional area. When combining *Ctnnb1-Venus* with *Cherry-Lef1* in the same mice, we observed that Cherry-Lef1 preceded junctional β-catenin-Venus expression in the RV (**Movie3**).

To track both the Cherry-Lef1 and β-catenin-Venus signals with higher resolution, we set up an adaptive feedback microscopy pipeline in which images acquired by the microscope are automatically processed by predefined image analysis routines (**Fig. 3a**). This allows for adjustment of image acquisition parameters and acquisition positions within the sample in real time. In this study, the adaptive feedback microscopy was used to collect high resolution images of individual nephrons over time. Live imaging with the feedback control system revealed that the Lef1 expression gradient is not static but appeared to be moving over time from the distal end towards the medial portion of the nephron (**Fig. 3b, c, d; Movie4**). While we could observe the movement of the Cherry-Lef1 peak in low resolution images (**Fig. 2d**), high resolution images were required for more precise analysis.

**Figure 3.**
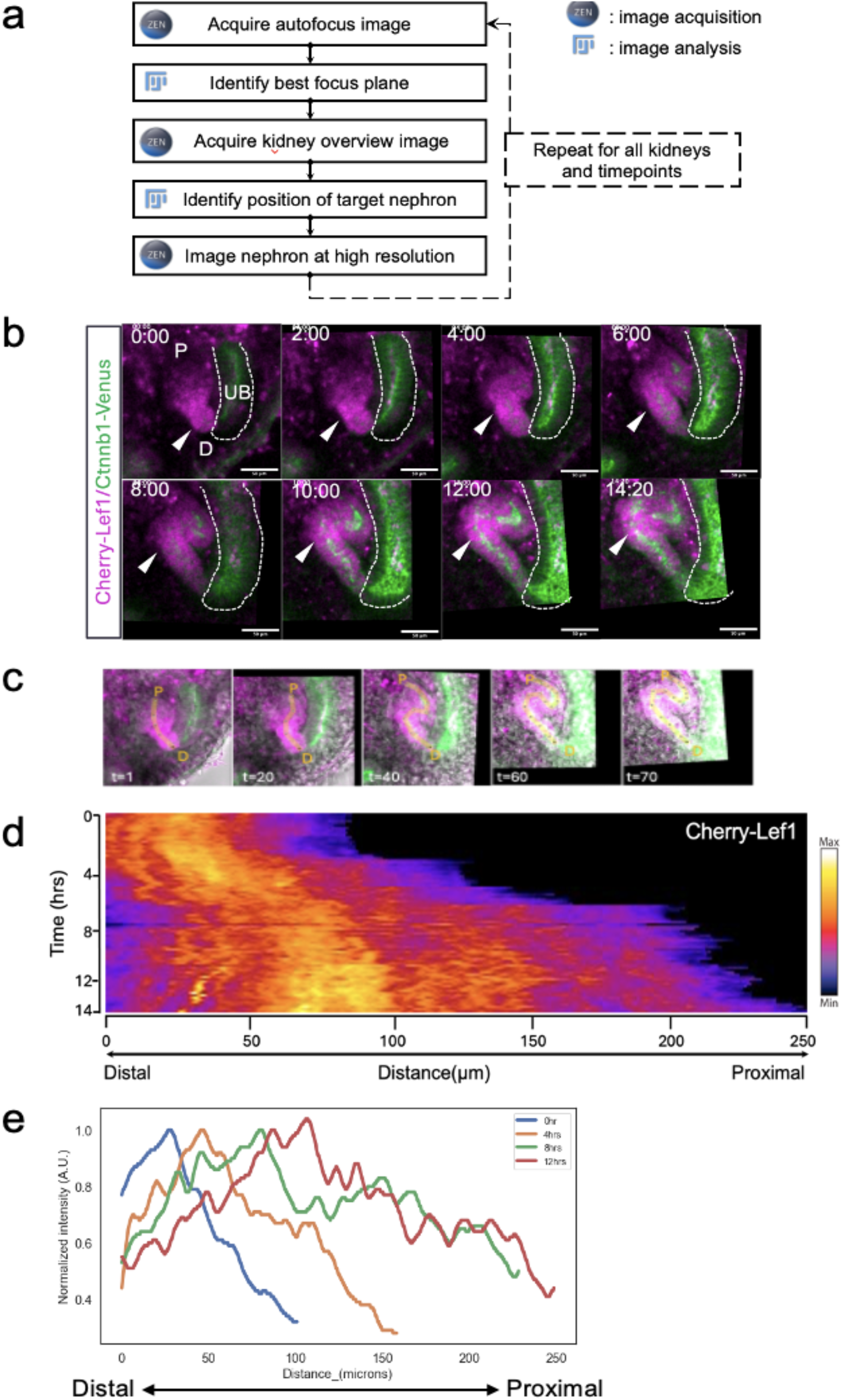
Real time imaging of Cherry-Lef1 and Ctnnb1-Venus protein signals in mouse nephron patterning by adaptive feedback control system. **(a)** Schematic representation of adaptive feedback control system. Microscope software ZEN interacts with Fiji to adjust image acquisition parameters in real time based on feedback from acquired imaging. **(b)** Snapshot images of real time imaging at different time points(0, 2, 4, 6, 8, 10,12 and 14.2hrs) in embryonic kidneys(E12.5) using Cherry-Lef1/Ctnnb1-Venus mouse line. Scale bars, 50µm. D: distal, P: proximal, UB: ureteric bud. The position of the highest intensity signal of Cherry-Lef1 is indicated by white arrow. **(c)** Orange lines depict ROI for kymograph and quantification of intensity at different time points(t). t=1: 10min, t=20: 200min, t=40: 400min, t=60: 600min, t=7: 700min. D: distal, P: proximal. **(d)** Intensity kymograph of Cherry-Lef1 signal in mouse nephron patterning. **(e)** Quantification of Cherry-Lef1 signal from distal to proximal at different time points (0,4,8 and 12hrs). Plot shows normalized intensity of Cherry-Lef1 signal from distal to proximal in nephron development.

Furthermore, to rule out that the Lef1 pattern is only seen in the *ex vivo* system, we isolated embryonic kidneys at E12.75 and conducted whole mount staining using an anti-Cherry antibody on E12.75 embryonic kidneys. The Cherry-Lef1 signal peak was observed at the medial part of the nephron in E12.75 embryonic kidneys (**Fig. 3 - Supplement 1a, b, c**).

Together, these data suggest that the Cherry-Lef1 expression gradient does not remain static but instead it is moving from distal to the proximal direction as the RV tissue undergoes lengthening and patterning.

## Methods

### Animal work

All animals were housed in the EMBL animal facility under veterinarians’ supervision and were treated following the guidelines of the European Commission, revised directive 2010/63/EU and AVMA guidelines 2007. All the animal experiments were approved by the EMBL Institutional Animal Care and Use Committee (project code: 21–001_HD_AA). The detection of a vaginal plug was designated as embryonic day (E) 0.5, and all experiments were conducted with E12.5, E12.75 or E13.5 embryos.

### Mouse lines

The mouse lines were generated in the Aulehla lab (EMBL) and their detailed characterization will be presented elsewhere. In brief: To generate *mCherry-linker-Lef1* knock-in alleles, we targeted the start codon of endogenous Lef1 locus with one selection and two different reporter cassettes coding for *mCherry linker* and *EGFP-linker*. The reporter cassettes were flanked by loxP- and FRT-sites to remove the selection cassette. Thus, the targeting vector was constructed as followed: *FRT-mCherry-linker-loxP-PGK-Neo-FRT-mEGFP-linker-loxP*. Cre-mediated excision resulted in the mCherry-linker-Lef1 allele used in this study.

To generate *Ctnnb1-linker-mVenus* knock-in alleles, we targeted the stop codon of the endogenous Ctnnb1 locus with one selection and two different reporter cassettes coding for linker mVenus and T2A-mCherry. The reporter cassettes were flanked by loxP- and FRT-sites to remove the selection cassette. Thus, the targeting vector was constructed as followed: l*oxP-linker-mVenus-FRT-PGK Neo-loxP-T2A-mCherry-FRT*. FRT-mediated excision resulted in *Ctnnb1-linker-mVenus* allele used in this study.

These mouse lines were generated by standard gene targeting techniques using R1 embryonic stem cells. Briefly, chimeric mice were obtained by C57BL/6 blastocyst injection and then outbred to establish the line through germline.

### Genotyping

*Cherry-Lef1* mice were genotyped by PCR using primers Lef1-wt-forward (CTTCCATTCACAGTCCTCCC), Lef1-Cherry-forward (AAGCAGAGGCTGAAGCTGAA), Lef1-wt-reverse (CTCCTCTTCGGGATGACTGA) produced bands: wild-type, 210-bp; Mutant, 434-bp. *Ctnnb1-Venus* mice were genotyped using Ctnnb1-wt-forward (GCAATCAGCTGGCCTGGTTT), Ctnnb1-Venus-forward (ATGGTCCTGCTGGAGTTCGT), Ctnnb1-wt-reverse (GCCTAAAACCATTCCCACCC) primers resulting in following bands: wild-type, 190-bp; Mutant, 230-bp.

### *Ex vivo* kidney culture

The embryonic kidneys were dissected from CD1 embryos crossed with *Ctnnb1-linker mVenus/mCherry-linker-Lef1* mouse line. 8-well chamber, Nunc™ Lab-Tek™ Chambered Coverglass (155411PK, Thermo Fischer) was coated with 160ul of Fibronectin (F1141-1MG, MERCK), which is diluted 1:20 with PBS per well, and incubated at RT for 2hrs. Fibronectin was removed from the chamber after 2hrs and equilibrated with 80ul of culture medium: DMEM/F12 (21041025, Thermo Fischer), 10% FBS, 1% penicillin/streptomycin for 30min. Dissected embryonic kidneys were cultured on fibronectin-coated chamber with 90ul of culture medium at 37°C and 5% CO_2_.

### Whole mount immunostaining

The embryonic kidneys were transferred to 4% PFA for 1 hr at 4°C, washed with PBST three times for 5min and stored at 4°C. For whole mount staining, samples were blocked with blocking buffer: 1% BSA, 0.2% nonfat dry milk powder, 0.3% Triton X-100 in PBS for 1 hour and incubated with Calbindin1 (1:100, 02401, Bicell Scientific), CDH1 (1:500, 610181, BD Bioscience), Six2 (1:100, 11562-1-AP, Proteintech), Cytokeratin-8 (1:50, MABT329, MERCK) and WT1 (1:1000, ab89901, Abcam) antibodies at 4°C overnight. Next day, samples were washed 3 times for 2 hours with PBST at RT and incubated with secondary antibodies: anti-mouse Alexa Fluor 488 (1:500, A11029, Thermo Fischer), anti-rabbit Alexa Fluor 568 (1:500, A10042, Thermo Fischer), anti-rat Alexa Fluor 633 (1:500, A21094, Thermo Fischer) for 1 hour at RT. After 3 times washes with PBST for 5min, samples were mounted with a drop of Vectashield mounting medium (H-1000-10, Vector Laboratories). Nikon AI microscopy was used for imaging.

### Confocal real time imaging and adaptive feedback control system

Imaging was performed on a Zeiss LSM780 NLO confocal microscope controlled by ZenBlack software (version 2.3 SP1). The chamber was placed into a chamber holder with wet tissue to keep humidity and inserted into an on-stage incubator at 37°C and 5% CO_2_. Samples were imaged with the Apochromat 20x/0.8 M27 air objective. β-catenin-Venus was excited with 488 nm laser and detected with 500-560 nm emission filter range. Cherry-Lef1 was excited with 561 nm laser and detected with 569-638 nm emission filter range. Multiple samples were recorded using a motorized stage during each experiment. Movies were recorded in pixel size, 1,4µm, image size, x: 708,49µm, y: 708,49µm, z:64µm in low zoomed image and in pixel size, 0,415µm, image size, x: 212,55µm, y: 212,55µm, z: 24µm in high zoomed image. The time-lapse images were captured every 10 min and an adaptive feedback microscopy pipeline was set up to track Cherry-Lef1 signal for high-resolution images.

Adaptive feedback microscopy was set up using a combination of MyPic macros for ZenBlack software (https://git.embl.de/grp-ellenberg/mypic), AutoMicTools library (https://git.embl.de/halavaty/AutoMicTools) for ImageJ/Fiji ^21,22^. The workflow consists of 3 image acquisition and 2 image analysis blocks. After moving the stage to the next kidney, the XZ scan in the reflection channel is first acquired and automatically processed to identify the position of a coverslip. This information is used to adjust objective focus position. Next, a low resolution image covering the entire kidney is acquired with a z-stack of 9 planes with 8 µm distance. This image is automatically processed to identify the actual position of a target nephron. This position is sent to the microscope to image this nephron at high resolution.

### Image and data processing

Fiji ^21^ was used for the analysis of the real time imaging data. Mouse nephron development movies were smoothened (gaussian blur, radius 5), a maximum intensity projection was made of slices containing tissue of interest and movies were registered using the ‘MultiStackReg’ function in Fiji. For quantification of the intensity of signal along the nephron axis from distal to proximal manual line ROIs were drawn for each time point (**Fig. 3c**). Intensity profiles along those lines were measured automatically and assembled into a Kymograph (**Fig. 3d**). The latter steps were implemented in a Fiji Groovy script which is available at https://git.embl.de/grp-cba/kidney-lif1-dynamics.

For quantification of mean intensity for whole mount staining images, images were smoothened (gaussian blur, radius 10), a maximal intensity projection was made by 10 z-slices containing nephron. Segmented line (line width, 20) was used to quantify the intensity of the nephron axis from distal to proximal.

## Discussion

Here, we employed a new method for renal organ culture that relies on direct plating of dissected embryonic kidneys on fibronectin-coated glass slides. While maintaining proper organ growth through the air-liquid-interface, this method overcomes limitations of the long working distance between the objective and the sample as well as the poor signal-to-noise ratios caused by the filter membrane. Furthermore, we applied the adaptive feedback microscopy to track Cherry-Lef1/β-catenin-Venus signal dynamics during nephron development at high resolution. The adaptive feedback microscopy workflow consists of several acquisition and image analysis blocks as described in Materials and Methods. Low resolution images of entire kidneys were automatically processed to identify positions of the tracked nephrons and image them at high resolution. The tools used here were applied in other studies to run adaptive feedback microscopy experiments ^23-26^. Moreover, we introduce two new Wnt reporter lines that allow for the visualization of the nascent nephron (*Cherry-Lef1* and *Ctnnb1-Venus*) and the ureteric bud (*Ctnnb1-Venus*). Since Cherry-Lef1 homozygous mice are viable, it indicates that the Cherry-Lef1 fusion protein is functional. Performing live imaging with these reporters, we could show that the Cherry-Lef1 signal forms a gradient in the distal to proximal direction during nephron development. Our feedback control imaging showed that this gradient does not seem to be static because its peak is moving into the medial-proximal direction.

While it can be assumed that the initial Lef1 expression is induced by Wnt9b ^3^, the question remains how far this morphogen can act. Our observation suggests that the Wnt9b effect is short-range and transient. This would be similar to the gut where Wnt3 that is expressed in the Paneth cells acts in a short-range manner with direct cell contact with the intestinal stem cell niche ^27^. In this manner, Frizzled-bound Wnt can spread passively by stem-cell division, which in turn dilutes surface-bound Wnt, thus creating a gradient. In the distal RV, a proliferative zone has indeed been observed that could account for a similar dilution of the Lef1 signal ^28^. Yet, it remains difficult to distinguish between a long-range gradient and a short-range gradient acting on proliferating target cells in the nephron. Therefore, it would be important to study tagged versions of Wnt9b to see how they interact with their target cells. Also, the role of cell proliferation in the RV in shaping the gradient and the Lef1 signal would need to be explored further ^28^.

Unlike the *Lef1* reporter, *Cnnb1-Venus* was not viable in homozygous condition, suggesting that the tag interferes with normal β-catenin function. Nuclear β-catenin was not detected, but needs to be confirmed with a nuclear marker. Nevertheless, it can be used as a marker for ureteric bud and RV adherens junctions. In the RV, Lef1 expression preceded β-catenin-Venus expression suggesting that β-catenin is not a component of the earliest adherens junctions in the developing nephron epithelium. Instead, these junctions might contain afadin and K-Cadherin as reported previously ^29,30^ that are then being replaced by β-catenin and its binding partner E-Cadherin during epithelial maturation.

In summary, we provide here a new imaging setup for visualizing Wnt signaling during nephron formation with unprecedented resolution. Follow-up studies will be needed to better understand the nature of the Wnt signaling gradient and the responses of the target cells within the developing nephron

## Acknowledgements

We thank Prof. Ryuichi Nishinakamura for providing advice on the project. We appreciate all the support of ALMF, LAR at EMBL and Nikon Imaging Center, Heidelberg. This work was supported by the Collaborative Research Center (SFB1324 on Wnt Signaling) funded by Deutsche Forschungsgemeinschaft SFB1324. We also acknowledge funding by the European Research Council (ERC) under the European Horizon 2020 research and innovation programme (Grant agreement No. 865408 (RENOPROTECT).

## Author Contributions

Designed the project: M.S., N.T.S., Performed experiments: N.T.S., Wrote the manuscript: N.T.S., S.H., M.S., Setup adaptive feedback control: A.H., Data analysis: N.T.S., S.H., C.T., A.H., Supervised the project: M.S.

---

**Figure S1 - Supplement 1.**
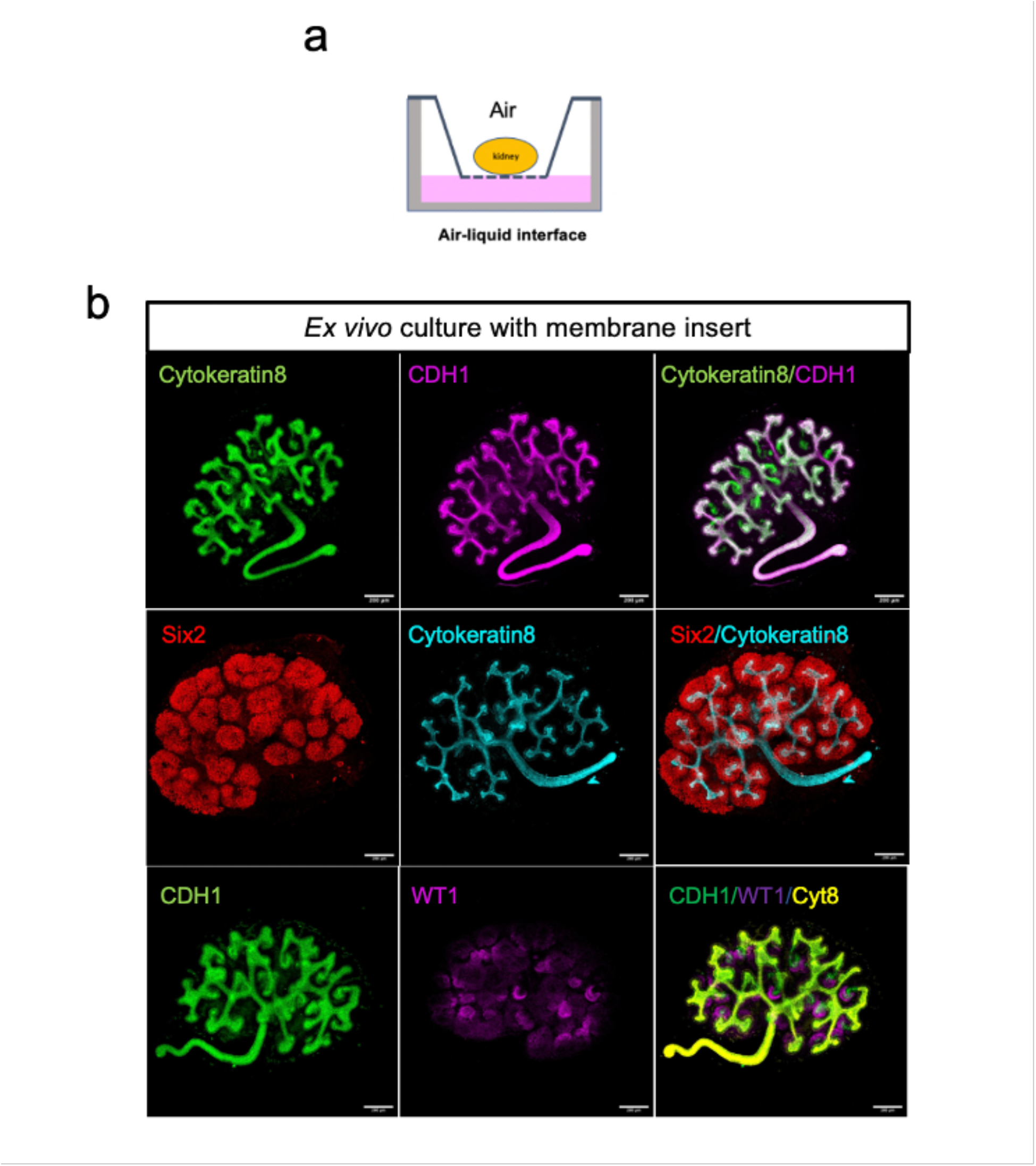
Development of mouse embryonic kidneys in traditional Trowell-type membrane insert culture. **(a)** Schematic of *ex vivo* culture with Trowell-type membrane insert. **(b)** Whole mount staining of embryonic kidneys after *ex vivo* culture for 18 hrs using Six2, Calbindin1, Cytokeratin8, CDH1 and WT1 antibodies. Scale bar, 200µm,

**Figure S2 - Supplement 1.**
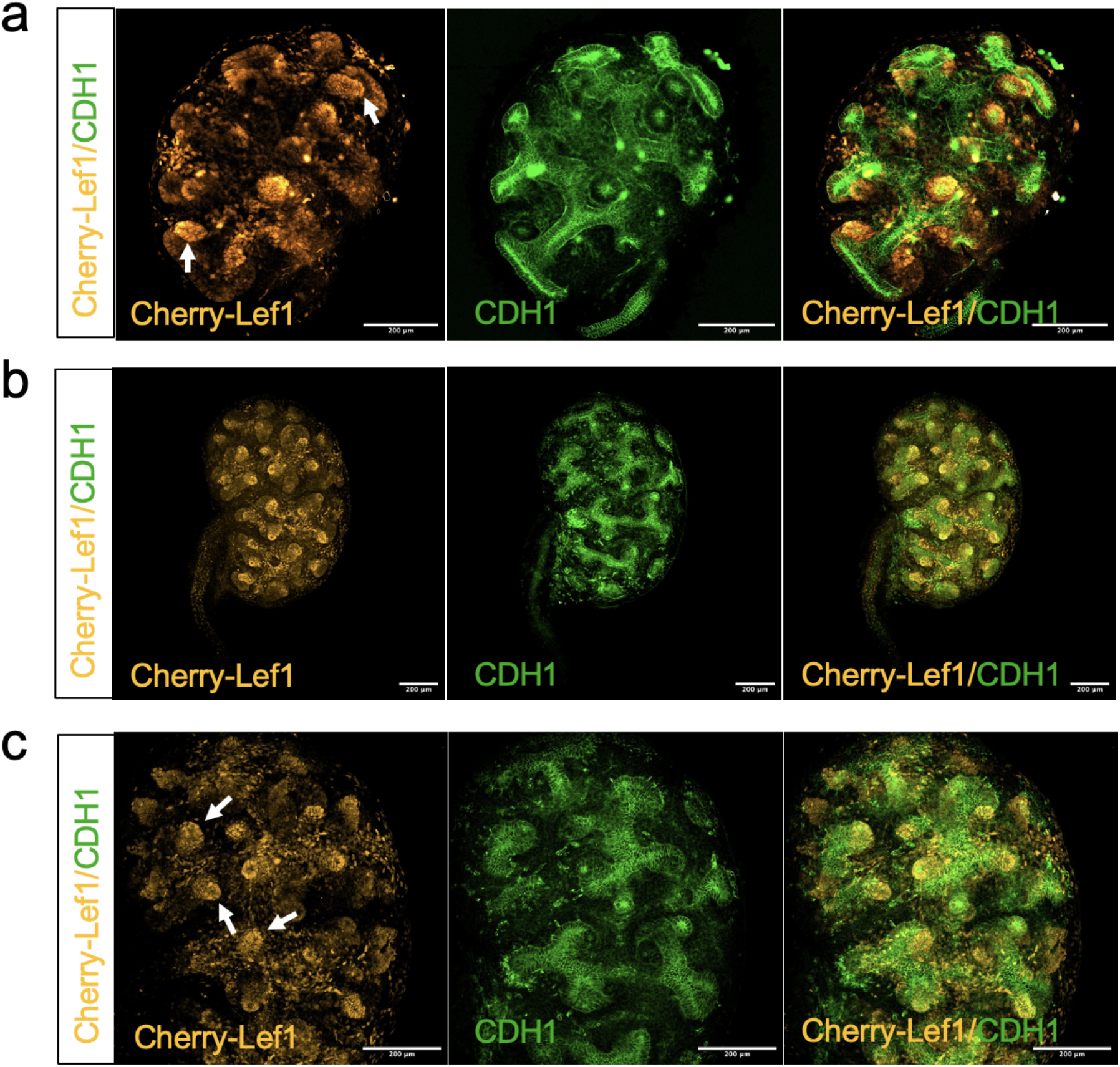
Cherry-Lef1 protein expression pattern in mouse embryonic kidney. **(a)** Whole mount staining of mouse embryonic kidney at E12.5 with Cherry and CDH1 antibodies. Scale bar, 200µm. White arrows show conical-shape like Cherry-Lef1 expression. **(b)** Whole mount staining of mouse embryonic kidney at E13.5 with Cherry and CDH1 antibodies. Scale bar, 200µm. **(c)** Enlarged images of (b) Scale bar, 200µm. White arrows show conical-shape like Cherry-Lef1 expression pattern at E13.5.

**Figure S3 - Supplement 1.**
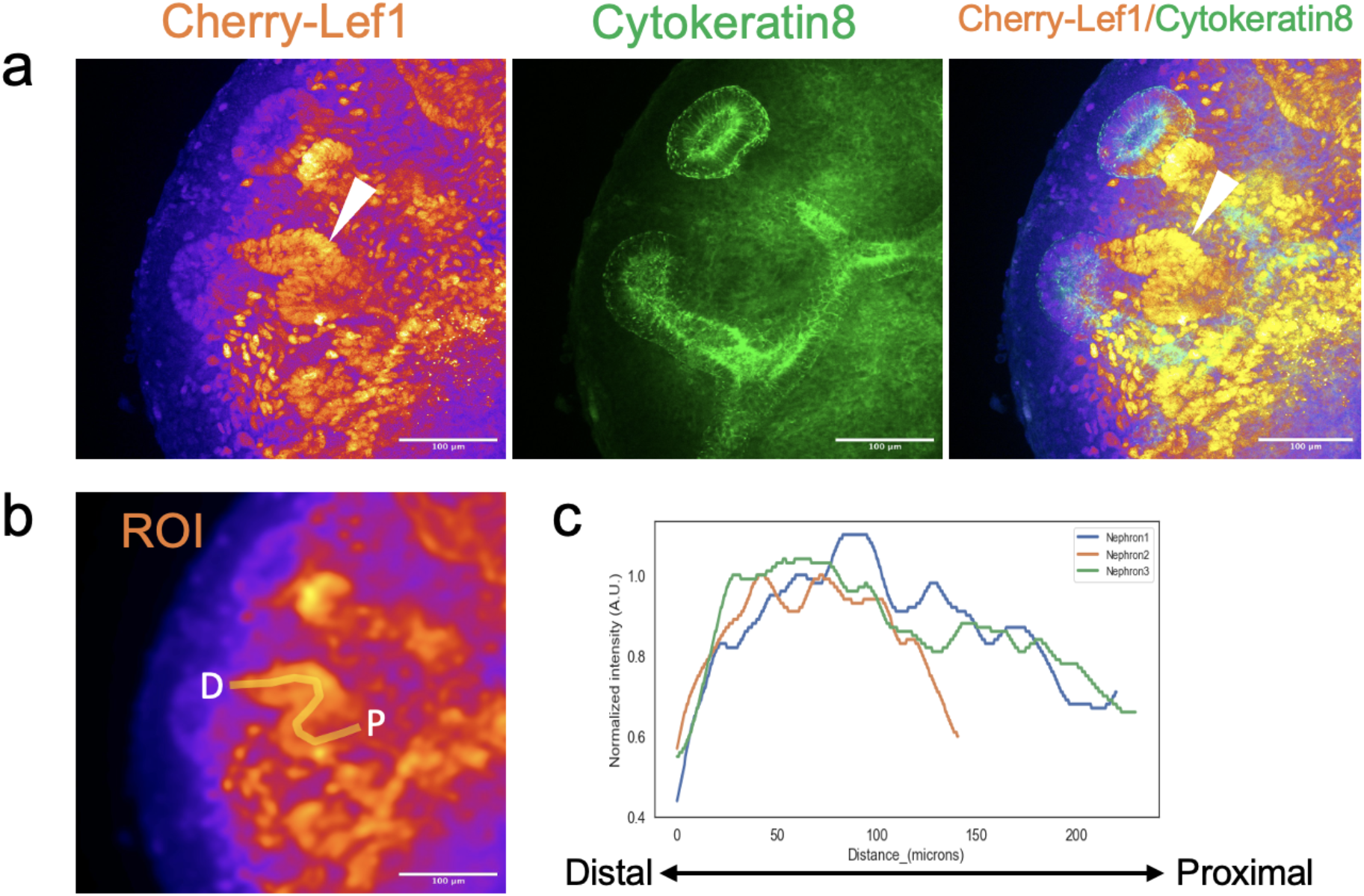
Cherry-Lef1 expression in mouse nephrons. **(a)** Whole mount staining of mouse embryonic kidney at E12.75 with Cherry and Cytokeratin 8 antibodies. White arrows indicate the highest signal of Cherry-Lef1 protein expression in the medial part of nephron. Scale bars, 100µm. **(b)** Roi was indicated as the orange line for quantification. D: distal, P: proximal. Scale bars, 100µm. **(c)** Quantification of normalized Cherry-Lef1 signal intensity from distal to proximal in mouse nephrons (n=3).

